# Increased reliability of visually-evoked activity in area V1 of the MECP2-duplication mouse model of autism

**DOI:** 10.1101/2022.02.27.482189

**Authors:** Ryan T. Ash, Ganna Palagina, Jiyoung Park, Jose A. Fernandez-Leon, Rob Seilheimer, Sangkyun Lee, Jasdeep Sabharwal, Fredy Reyes, Jing Wang, Dylan Lu, Sam Wu, Stelios M. Smirnakis

**Author notes:** These authors contributed equally to this work (Ganna Palagina, Jiyoung Park, Jose A. Fernandez-Leon). **Address correspondence to:** Stelios M. Smirnakis, MD, Ph.D., Brigham and Women’s Hospital, Rm #328, Department of Neurology, Harvard Medical School, 75 Francis Street, Boston, MA 02115.

## Abstract

Atypical sensory processing is now thought to be a core feature of the autism spectrum. Influential theories have proposed that both increased and decreased neural response reliability within sensory systems could underlie altered sensory processing in autism. Here, we report evidence for abnormally increased reliability of visual-evoked responses in layer 2/3 neurons of adult primary visual cortex in the MECP2-duplication syndrome animal model of autism. Increased response reliability was due in part to decreased response amplitude, decreased fluctuations in endogenous activity, and decreased neuronal coupling to endogenous activity. Similarly to what was observed neuronally, the optokinetic reflex occurred more reliably at low contrasts in mutant mice compared to controls. Retinal responses did not explain our observations. These data suggest that the circuit mechanisms for convolution of sensory-evoked and endogenous signal and noise may be altered in this form of syndromic autism.

## INTRODUCTION

Autism spectrum disorder (ASD) is classically a disorder of repetitive behaviors and impaired communication, but it is becoming better recognized that sensory perceptual dysfunction is a core feature of the disorder and is now a DSM5 ASD diagnostic criterion (Simmons et al., 2009; Grzadzinski et al., 2013; Robertson and Baron-Cohen, 2017). The great majority of individuals with ASD report hyper-or hyposensitivities in multiple sensory domains (Grzadzinski et al., 2013). In psychophysical studies, ASD subjects can show remarkable impairment or enhancement in perceptual sensitivity depending on task parameters. For example, autistic individuals demonstrate accelerated visual search, enhanced motion perceptual sensitivity under low stimulus noise, and decreased susceptibility to visual illusions, while also showing impaired recognition for faces and biological motion, decreased motion perception under high stimulus noise, and visual sensory hypersensitivity (Simmons et al., 2009; Robertson and Baron-Cohen, 2017). Neurophysiological responses in the visual system are altered in autistic individuals (McCleery et al., 2007; Isler et al., 2010; Dinstein et al., 2012; Pei et al., 2012; Weinger et al., 2014; Takarae et al., 2016).

Recent influential theories propose that disrupted neuronal response reliability underlies sensory processing deficits in autism (Dinstein et al., 2012; Haigh et al., 2015). While some have theorized that *excess* variability of endogenously generated activity (assessed either behaviorally or neurophysiologically) underlie abnormal reliability in autism (Simmons et al., 2009; Heeger et al., 2017; Park et al., 2017), others have proposed that there is *insufficient* endogenous variability in autism (Brock, 2012; Pellicano and Burr, 2012; Davis and Plaisted-Grant, 2015). Significant decreases in neuronal response reliability have been observed in ASD patients (Milne, 2011; Dinstein et al., 2012; Weinger et al., 2014; Haigh et al., 2015, 2016, 2020; Otto-Meyer et al., 2018) and animal models of autism (Banerjee et al., 2016; Geramita et al., 2020), but well-controlled studies have also found evidence for the opposite pattern (Coskun et al., 2009; Frey et al., 2013; Greenaway et al., 2013; Butler et al., 2017). Little work has been done to measure sensory-evoked and endogenous variability in cortical neurons in animal models of autism.

Here we measure response reliability in the mouse model of *MECP2* duplication syndrome (Collins et al., 2004; Ramocki et al., 2010), a highly-penetrant syndromic ASD caused by genomic duplication of methyl-CpG-binding protein 2 (Ramocki et al., 2010). This animal model demonstrates a range of autism-associated behaviors including repetitive stereotyped behaviors, abnormal ultrasonic vocalizations, and social avoidance as well as atypical sensory processing (Collins et al., 2004; Samaco et al., 2012; Jiang et al., 2013; Sztainberg et al., 2015; Zhang et al., 2017; Zhou et al., 2019; Ash et al., 2021). We find that response reliability is abnormally increased in visual cortical neurons in this mouse model. We identify three potential mechanisms for increased response reliability: decreased sensory-evoked responses, decreased fluctuations in endogenous spontaneous activity, and decreased coupling of neurons to endogenous fluctuations (Okun et al., 2015; Lee et al., 2019). Enhanced reliability at low contrasts was associated with enhanced optokinetic reflex responsiveness in these animals suggesting that changes in response reliability may propagate to midbrain circuits and influence perception. These data weigh against recent proposals for increased neural variability as a canonical pathophysiological mechanism in autism (Heeger et al., 2017), and instead support theories for insufficient endogenous noise in ASD (Davis and Plaisted-Grant, 2015).

## RESULTS

### Contrast-response functions are abnormal in the primary visual cortex of MECP2-duplication mice

Neuronal responses to oriented gratings were recorded for a range of contrasts by in vivo 2-photon calcium imaging in GCaMP6s-expressing (Chen et al., 2013) L2/3 neurons in area V1 of F1 C57;FVB adult (4-6-month-old) MECP2-duplication mice (Tg1, see Collins et al., 2004) and control littermates under fentanyl-dexmetotomidine sedation (note that Tg1 mice on F1 C57;FVB background do not have seizures). Visual stimuli consisted of oriented gratings at 12 different directions (0, 30…330°) and 3 contrasts (10, 30, 100%), randomly-interleaved (**Fig. 1A,** see Methods). We recorded from 7 transgenic mice and 4 control littermates, identifying 238 and 223 layer 2/3 neurons, respectively. The percentage of neurons that were visually responsive (see Methods) was similar between control and mutants (46±19%, 102 cells, and 51±12%, 114 cells, respectively). The rest of the analysis was restricted to these visually responsive cells.

**Figure 1.**
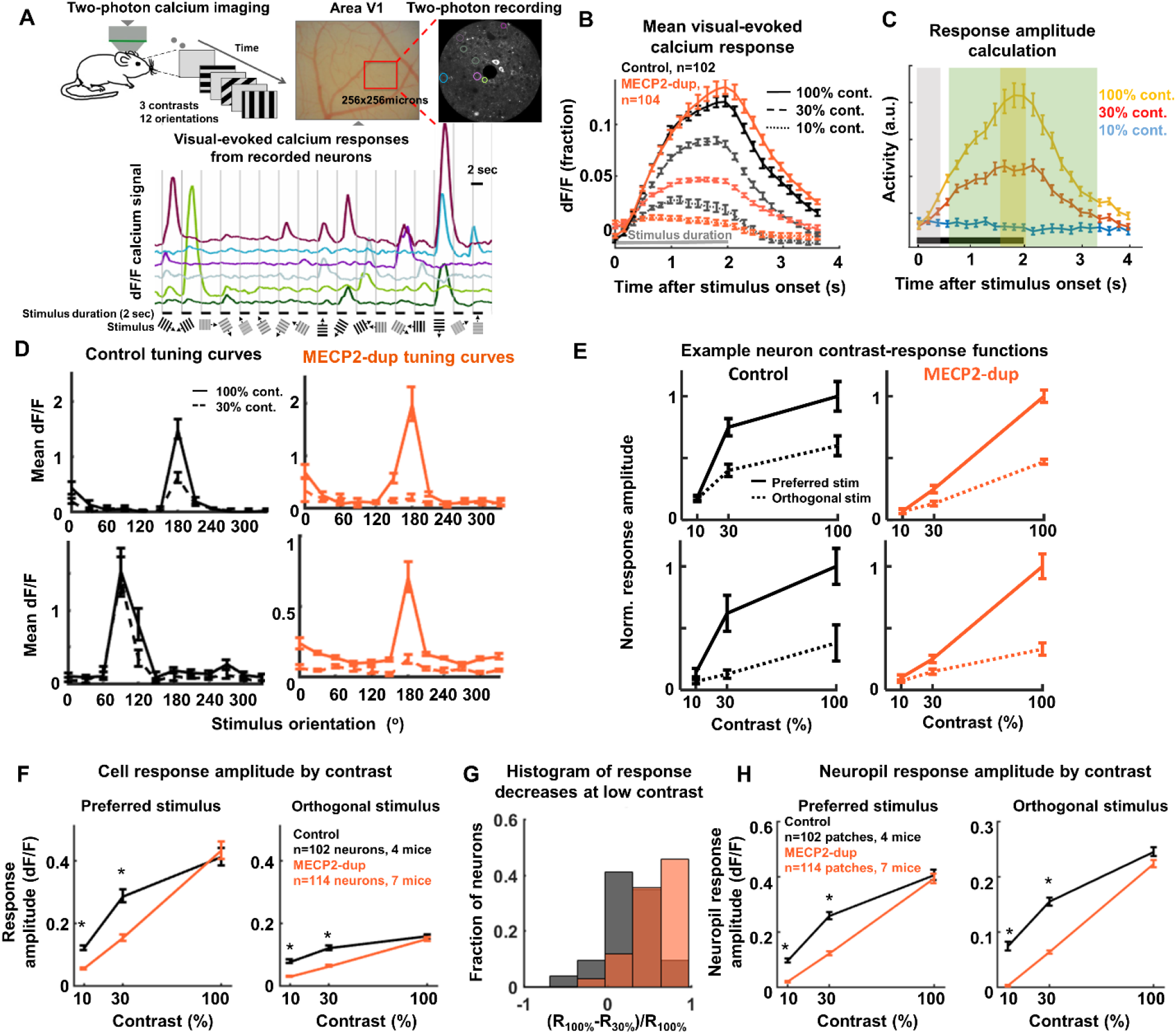
Abnormal contrast-response functions in primary visual cortex of MECP2-duplication mice. **A.** *Top left,* Experiment schematic. Mice were head-fixed under the 2-photon microscope, and randomly-interleaved oriented gratings (12 directions, 3 contrasts) were presented. *Top right,* sample optical image of a cranial window and averaged 2-photon movie showing V1 neurons expressing GCaMP6s. *Bottom,* example fractional dF/F calcium responses to oriented gratings. Tracing color depicts the recorded neuron circled in top right panel. Vertical lines depict stimulus onset. Horizontal lines, stimulus duration (2 sec). Grating images show the stimulus on each trial. Intertrial period was 2 sec. **B.** Visual-evoked calcium transients at different contrasts. Transients are computed by averaging across all trials, across all orientations, per neuron, then averaging across neurons. Error bars calculated across neurons; Control: n=102 neurons from 4 mice; MECP2-duplication, n=114 neurons from 7 mice. **C**. Illustration of how per-trial responses are calculated. Average responses of one neuron to the three contrasts are shown as yellow, red, and blue error bars. The response for each trial is calculated as the average of the 3 frames centered at the peak of the mean 100%-contrast response across all orientations +/- 1 frame (yellow bar). The peak is detected within the frames denoted by the green box. The black line shows the stimulus duration. The grey box shows the frames in which pre-stimulus activity is calculated (see methods). **D.** Example tuning curves from control (*left*) and *MECP2*-duplication (*right*) neurons at 100% (solid) and 30% (dashed) contrasts. These neurons did not display noticeable tuning at 10% contrast. **E.** Example single-neuron contrast-response functions, at the preferred (solid) and non-preferred (dotted) stimulus, normalized to the response at 100% contrast. Black: control. Orange: MECP2-duplication mice. Note that control neuronal responses increase at low contrasts then begin to saturate, while mutant responses increase more linearly with contrast. **F.** Response amplitude as a function of contrast. *<u>Left panel</u>:* dF/F response amplitude to each neuron’s preferred stimulus (peak of the orientation tuning curve) averaged across neurons as a function of contrast. *<u>Right panel</u>:* Similar plot for the *non-preferred* stimulus, i.e. gratings of orthogonal orientation to the peak of the tuning curve. **G**, Histogram of the fractional decrease of the response amplitude to the preferred stimulus from 100% to 30% contrast ((R_100%_-R_30%_)/R_100%_), across neurons. A value of 1 indicates complete loss of responsiveness at 30% contrast, while −1 indicates doubling of response amplitude at 30% contrast. Gray: control; Orange: MECP2-duplication. **H**. Neuropil response amplitude by contrast. * P<0.01, 2-way ANOVA followed by Tukey test for multiple comparisons. All values are shown as mean±SEM.

Spontaneous activity levels were first assessed by quantifying the rates of calcium events while the fentanyl-sedated mice viewed the mean luminance background. DF/F calcium traces were thresholded (see Methods) (Lu et al., 2016), and peaks were counted. Spontaneous event rates were slightly decreased in MECP2-duplication mice (0.027±0.001 Hz, n=114) compared to controls (0.033±0.002 Hz, n=102, P=0.04, 2-way ANOVA followed by Tukey post-hoc test), and spontaneous calcium transient amplitudes trended towards a slight decrease in mutant animals (MECP2-dup: 1.8±0.05 dF/F, control: 2±0.1 dF/F, P=0.06).

Visually-evoked responses averaged across all orientations and neurons (**Fig. 1B**) showed a strong interaction between genotype and contrast: Neuronal responses to low-contrast stimuli (dashed line 30%, dotted line 10%) were much decreased, whereas responses to high (100%, solid line) contrast stimuli were similar to controls, if not somewhat higher, in the mutant (**Fig. 1B**).

Response amplitudes were calculated for each visually responsive neuron across trials (**Fig.1C**) to generate orientation tuning curves (**Fig. 1D**). The response to the preferred stimulus markedly decreased at low contrasts in the majority of MECP2-duplication L2/3 V1 cells (**Fig. 1D**). **Fig. 1E** shows example contrast-response functions for control (left) and mutant (right) neurons. Contrast-response curves obtained from the preferred stimulus (solid lines) show the same functional trend (**Fig. 1E**) as curves obtained from the orthogonal stimulus (dotted lines); curves are normalized to the response of the high contrast preferred stimulus. While the response of the majority of control neurons increases significantly from 10% to 30% contrast then levels off, mutant neurons increase their response amplitude approximately linearly with contrast. Plotting the mean response amplitude across visually responsive neurons to the preferred and non-preferred stimulus (**Fig. 1F**) revealed a dramatic decrease in V1 neuronal responses specifically at low contrasts for mutant mice compared to controls (P < 0.01; Tukey post-hoc pairwise comparison following 2-way ANOVA; n=102 control, 114 MECP2-duplication neurons). The difference between mutants and controls remains significant also across animals (P=0.02 at 30% contrast, n=7 MECP2-duplication mice, 4 control; responses are normalized to the high-contrast preferred stimulus response per cell). **Fig. 1G** histograms the fractional decrease in response amplitude between 100% and 30% contrast ((R_100%_-R_30%_)/R_100%_) demonstrating that the responses of mutant neurons drop markedly as contrast decreases (**Fig. 1G**).

The decrease in response amplitude at low contrasts was also clearly observable in calcium signals arising in the neuropil surrounding cells (**Fig. 1H**). The neuropil signal arises from calcium transients in dendrites and axons arising from hundreds to thousands of GCaMP6s-expressing neurons whose processes lie within or traverse L2/3 of area V1. The neuropil signal gives a good average readout of overall V1 population activity and is known to correlate well with the local field potential (LFP) (Kerr et al., 2005). Decreased neuropil responses therefore indicate decreased aggregate evoked activity in primary visual cortex during low-contrast visual stimulation in the MECP2-duplication mouse.

Orientation selectivity index (OSI= (R_preferred_-R_orthogonal_)/R_preferred_) (Mazurek et al., 2014)) was similar between genotypes at 100% contrast (control: 0.32±0.02; MECP2-dup: 0.35±0.02, P=0.05, Tukey post-hoc pairwise comparison following 2-way ANOVA), while at 30% contrast OSI decreased in mutants (0.2±0.015 at 30% contrast versus 0.35±0.02 at 100% contrast, P<0.01) but remained approximately unchanged in controls (0.27±0.02 at 30% versus 0.32±0.02 at 100% contrast). At 10% contrast, the responses were weak and OSI calculation was not reliable (control: 0.12±0.01, MECP2-dup: 0.08±0.01). In agreement with the decreased response amplitude and fall in orientation selectivity at 30% contrast, the tuning width of mutant neurons increased at 30% contrast (64±2°) compared to control (54±2°; P<0.01, Tukey post-hoc pairwise comparison following 2-way ANOVA). At 100% contrast tuning widths remained similar (control: 53±2°, MECP2-dup: 58±2°). Direction selectivity in MECP2-duplication mice behaved similarly to orientation selectivity, decreasing significantly for stimuli at 30% contrast (control: 0.13±0.01, MECP2-dup: 0.073±0.006; p<0.01, Tukey pairwise comparison after 2-way ANOVA) but remaining essentially unchanged at 100% contrast (control: 0.17±0.01, MECP2-dup: 0.14±0.01; p > 0.05).

### Response Reliability as a Function of Contrast in V1 neurons of MECP2 duplication mice

As expected (Werner and Mountcastle, 1963), neuronal responses in our experiment fluctuated from trial to trial, even in response to the same stimulus (**Fig. 2A**). We quantified the variation in each neuron’s response amplitude by measuring the Fano factor (FF, defined as response variance divided by response mean, σ^2^/μ) across multiple repetitions of the same stimulus (**Fig. 2B**). Interestingly, we observed a significant *increase* in response reliability (decreased FF) in MECP2-duplication compared to control neurons (compare **Fig. 2A** top row to bottom row) that, similar to the response amplitude changes, was more pronounced to low (10% and 30%) contrast stimuli (**Fig. 2C**). The effect was observable in responses to both the preferred (left panel, P<0.01 at 10% contrast, P<0.01 at 30% contrast, P=0.09 at 100% contrast, Tukey post-hoc pairwise comparison following 2-way ANOVA, n=102 control neurons, 114 MECP2-dup neurons) and orthogonal (right panel, P<0.01 at 10%, P<0.01 at 30%, P=0.39 at 100% contrast) stimuli. Although FF decreases with decreasing neuronal response amplitude (**Fig. 2D**) as expected (Goris et al., 2014), the decrease in response variability in mutants could not be explained simply by the decrease in firing at low contrasts, as the difference in FF was observable across a range of mean response amplitudes (**Fig. 2D**). Note also that our observations cannot be explained by non-specific decreased variability in the calcium responses of mutant animals, because they are apparent in the relative magnitude of the Fano Factors between 100% and 30% contrast, and results were the same whether they were measured as calcium Fano Factors or spiking Fano factors (i.e. by dividing each Fano factor by the Fano factor at 100% contrast to factor out the calcium-related response constant, see Montijn et al., 2014).

**Fig. 2.**
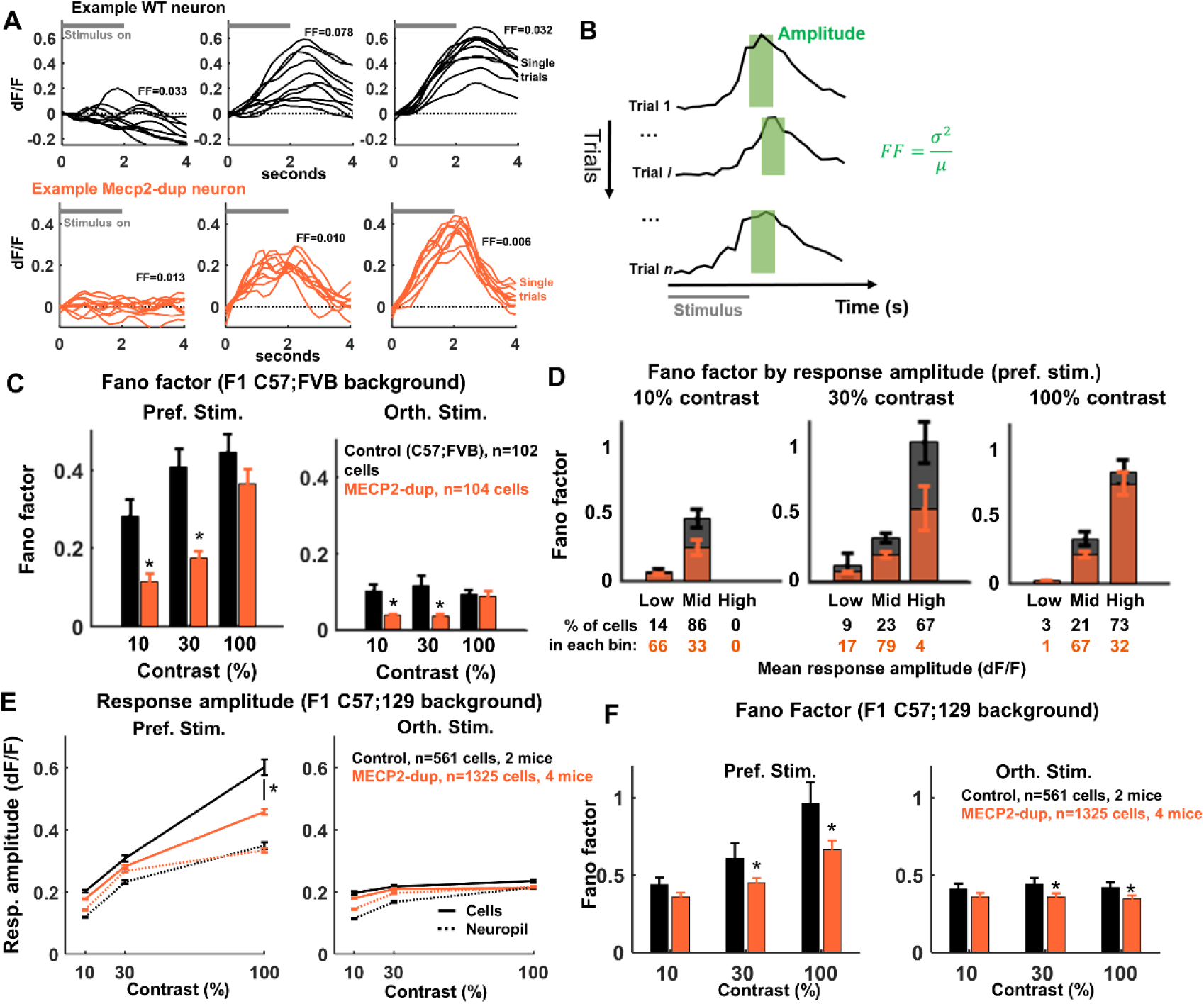
Response variability of V1 neurons drops at low contrast in MECP2-duplication mice. **A.** Single trial responses recorded from an example WT neuron (top row) and MECP2-duplication neuron (bottom row) across three different contrasts (10%, 30%, and 100% left to right). Each trial response was baseline subtracted and smoothed with a 5-bin moving averages filter for illustration putposes. Horizontal grey line in top left of each paneldepicts stimulus on time. Calculated Fano Factor is indicated for each set of responses. **B**. Schematic showing how the Fano Factor (FF=variance/mean) across trials was calculated. **C.** FF of calcium responses as a function of stimulus contrast for responses to the preferred (left) and the orthogonal (right) stimulus, across all visually responsive neurons, in F1 C57;FVB genetic background mice. *Control:* n=102 neurons from 4 mice; *MECP2-duplication:* n=114 neurons from 7 mice. Note that response variability of V1 neurons drops at low contrast. The effect remained statistically significant (P<0.05 at 10% and 30% contrast) when FF was calculated per-animal (Tukey pairwise comparison following 2-way ANOVA). **D**. FF binned by response amplitude to the preferred stimulus at each contrast. Neurons were grouped into bins of low (0-0.34), moderate (0.34-1.35) and high (1.35-5.3) activity. Values below each bar denote the percentage of cells in each bin. Note that the Fano factor decreases with decreasing neuronal response amplitude, but differences in mutants vs. controls cannot be explained simply as a result of different firing rates elicited by different contrasts. **E, D**. Replication of decreased V1 response variability in an independent genetic background and calcium imaging setup. The replication experiment was identical to the main experiment except in two regards: Experimental animals were F1 C57;129 (vs. F1 C57;FVB), and GCaMP6s was expressed transgenically in layer 2/3 pyramidal neurons under the thy1 promoter (vs. pan-neuronal viral expression using injection of AAV-GCaMP6s), allowing more cells recorded per mouse and ensuring that all recorded neurons are pyramidal cells. **E**. Response amplitude in F1 C57;129 mice for cells (solid lines) and surrounding neuropil (dotted lines) for the preferred (left panel) and non-preferred (right panel) stimulus. *Control:* n=561 neurons from 2 mice; *MECP2-duplication:* n=1325 neurons from 4 mice. Error bars depict 99% confidence intervals. **F**. Fano Factor (response variability) in F1 C57;129 mice for the preferred (*left panel*) and orthogonal (*right panel*) stimulus, across all visually responsive neurons, showing a replication of decreased response variability in MECP2-duplication mice.

### Reported changes in response reliability are not genetic background-specific

We wanted to replicate our finding of increased response reliability in an independent data set with a different genetic background. 129-background MECP2-duplication mice were crossed to C57 thy1-GCaMP6 mice to generate F1 hybrid C57;129 experimental animals, which genomically express GCaMP6 in excitatory neurons (in contrast to the previous experiment which used C57;FVB animals and virally-mediated GCaMP6 expression), allowing ensured specificity to pyramidal neurons (Thy1-GCaMP6 is pyramidal-neuron specific, while with virally-mediated GCAMP6 a subset of cells will be GABAergic) and more cells recorded per mouse. In the replication experiment, similar to what was observed in F1 C57;FVB mice, V1 neuronal response amplitude was decreased in mutants vs. controls, this time more pronouncedly in responses to high-contrast stimuli (**Fig. 2E**). Response variability (FF) was also decreased, this time observable across all contrasts and for both preferred and non-preferred visual stimuli (**Fig. 2F**). We note that there were differences in the measured values of response amplitude and Fano Factor in the replication experiment compared to the original experiment, likely due to a combination of differences in genetic background, GCaMP expression method, and microscope used, which do not affect the validity of the comparisons between mutant animals and littermate controls.

### Potential mechanisms for increased response reliability in MECP2-duplication mice

We hypothesized that the decreased neuronal response variability at low contrasts could be due to a decrease in cortical activity fluctuations, and/or a decreased coupling of V1 neurons to these cortical fluctuations (Ecker et al., 2016; Denfield et al., 2017). Neuropil activity reflects the aggregate of calcium transients in axons and dendrites immediately surrounding the soma and thereby can serves as a measure of cortical activity flux (Grinvald, 2005; Kerr et al., 2005). Neuropil visual responses were much weaker in mutants compared to controls (**Fig. 1H**) at low contrast, and neuropil response variability (FF) across trials was also significantly decreased (**Fig. 3A**). These observations suggest that global, trial-to-trial, fluctuations in activity are significantly decreased in mutant mice when low-contrast stimuli are presented. In addition, V1 neurons appear to be more weakly coupled to aggregate activity in mutant animals: On average the Pearson correlation coefficient between the activity of a neuron and the surrounding neuropil was significantly decreased in MeCP2-duplication animals, particularly at low contrasts (**Fig. 3B**). Note that Pearson correlation was computed *after* subtracting the estimated optical neuropil contamination from the somatic signal (see Methods, (Lee et al., 2017)), so correlation values reported here are unlikely to reflect significant optical neuropil contamination. In any event, even if there were a degree of residual optical contamination this would not explain our results, as there was no significant difference in optical quality in the mutant versus the control preparations. Note also that no thresholding was performed that might decrease variability or artificially bias cell-neuropil correlations by setting small responses to zero.

**Fig. 3.**
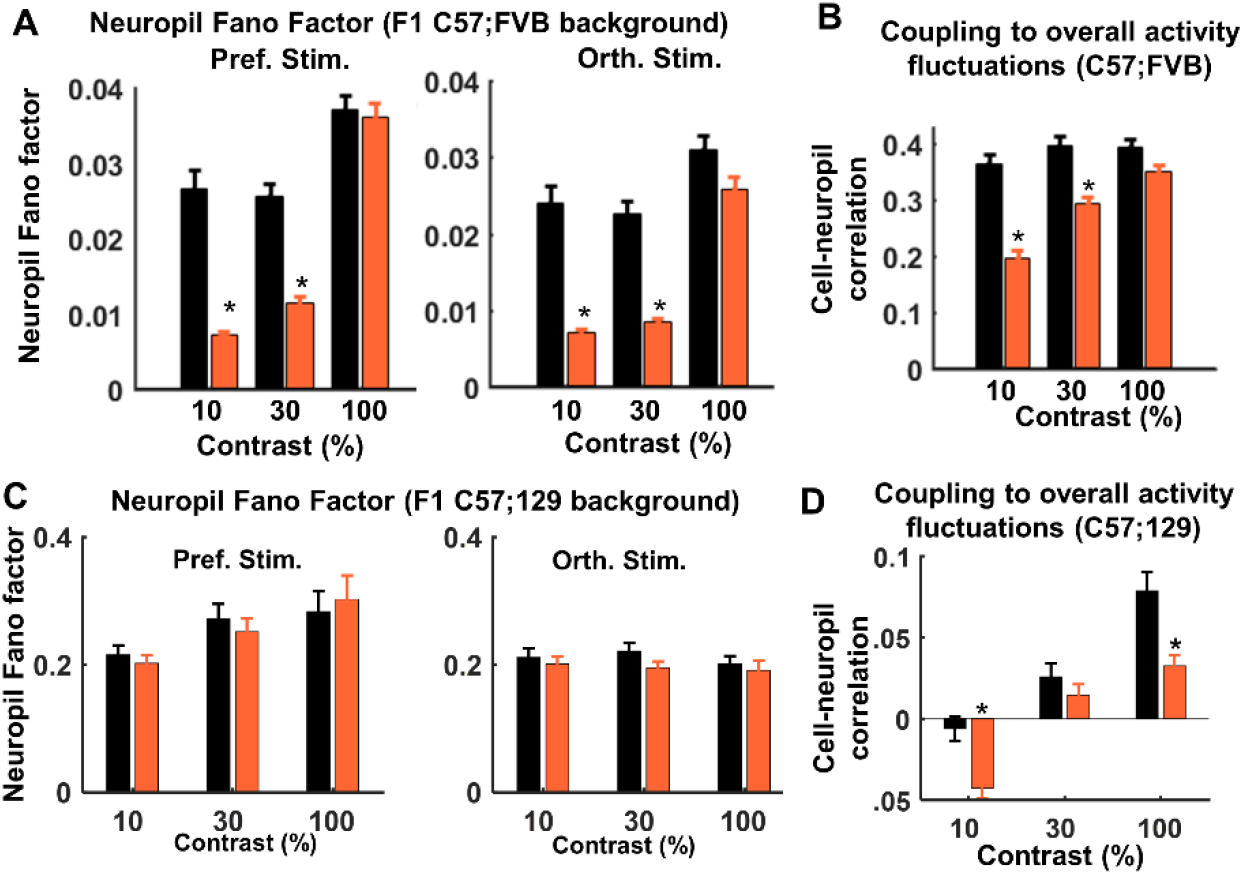
Potential mechanisms for decreased response variability in MECP2-duplication mice. **A.** Neuropil FF by contrast in C57;FVB background mice for preferred (left panel) and orthogonal non-preferred (right panel) stimuli. *Control:* n=102 neurons from 4 mice; *MECP2-duplication:* n=114 neurons from 7 mice. **B**. Coupling of each neuron to aggregate evoked visual cortical activity fluctuations (neuropil activity). The correlation coefficient between each cell’s responses and the activity of the surrounding neuropil, after correcting for optical contamination (see Methods), averaged across cells. * P<0.01, Tukey post-hoc pairwise comparison following 2-way ANOVA. All values are shown as mean±SEM. **C**. Neuropil Fano Factor, as in panel A, in the replication experiment with F1 C57;129 mice and transgenic thy1-GCaMP6 expression. *Control:* n=561 neurons from 2 mice; *MECP2-duplication:* n=1325 neurons from 4 mice. **D**. The correlation coefficient between each cell’s responses and the activity of the surrounding neuropil, averaged across cells, in C57l129 background mice. * P<0.01, Tukey post-hoc pairwise comparison following 2-way ANOVA.

Interestingly, neuropil response variability was not decreased in mutant mice in the C57;129 background (**Fig. 3C**), suggesting that spontaneous fluctuations in background activity are not as affected. Nonetheless, the coupling to background fluctuations was signficinatly decreased in mutants vs. WT in the C57;129 genetic background as well (**Fig. 3D**).

These results reveal that visual response variability is significantly decreased in MECP2-duplication animals across two different genetic backgrounds. Furthermore, this is driven in part by the reduced coupling of individual layer 2/3 V1 neurons to aggregate evoked neuropil fluctuations, which correlates with moment-to-moment activity in incoming inputs (Kerr et al., 2005).

### Retinal responses in MECP2-duplication mice do not explain our findings

We wanted to investigate whether abnormal contrast-response functions in MECP2-duplication V1 neurons could be due to abnormal retinal input. Electroretinogram responses to full field flash stimuli at different luminance intensity, recorded from the optic nerves of 4 MECP2-duplication mice and 4 littermate controls *in vivo*, revealed no difference in the amplitude of the a or b waves between genotypes (**Fig. 4A-C**), suggesting no major functional abnormality in the photoreceptor or inner retinal layer. Interestingly, *ex vivo* retinal ganglion cell (RGC) responses from mutants were similar at low contrasts and actually increased at high contrasts relative to control responses (**Fig. 4D-E**, P=0.0001, t=-12.2, genotype x contrast interaction; P=0.02, t=2.2, effect of genotype; P=0.0005, t=3.4, effect of contrast; n=55 control cells, 97 MECP2-dup cells, linear mixed-effects models ANOVA, neurons nested within retinae). Importantly, the contrast at which we observe a prominent decrease in the firing of MECP2-duplication V1 neurons (30%) generates essentially normal if not slightly higher (non-significant trend) RGC responses (**Fig. 4E**). Similarly, Ex vivo retinal ganglion cell Fano Factors were not significantly different between genotypes (10% contrast, MECP2-dup: 2.6± 0.3, control: 3.3±0.5; 30% contrast, MECP2-dup: 2.2±0.2, control: 2.7±0.4; 100% contrast, MECP2-dup: 2.3±0.3, control: 2.3±0.4). These data suggest that changes in retinal function are not likely to account for the decreased responsiveness and variability observed in mutant cortical neurons.

**Figure 4.**
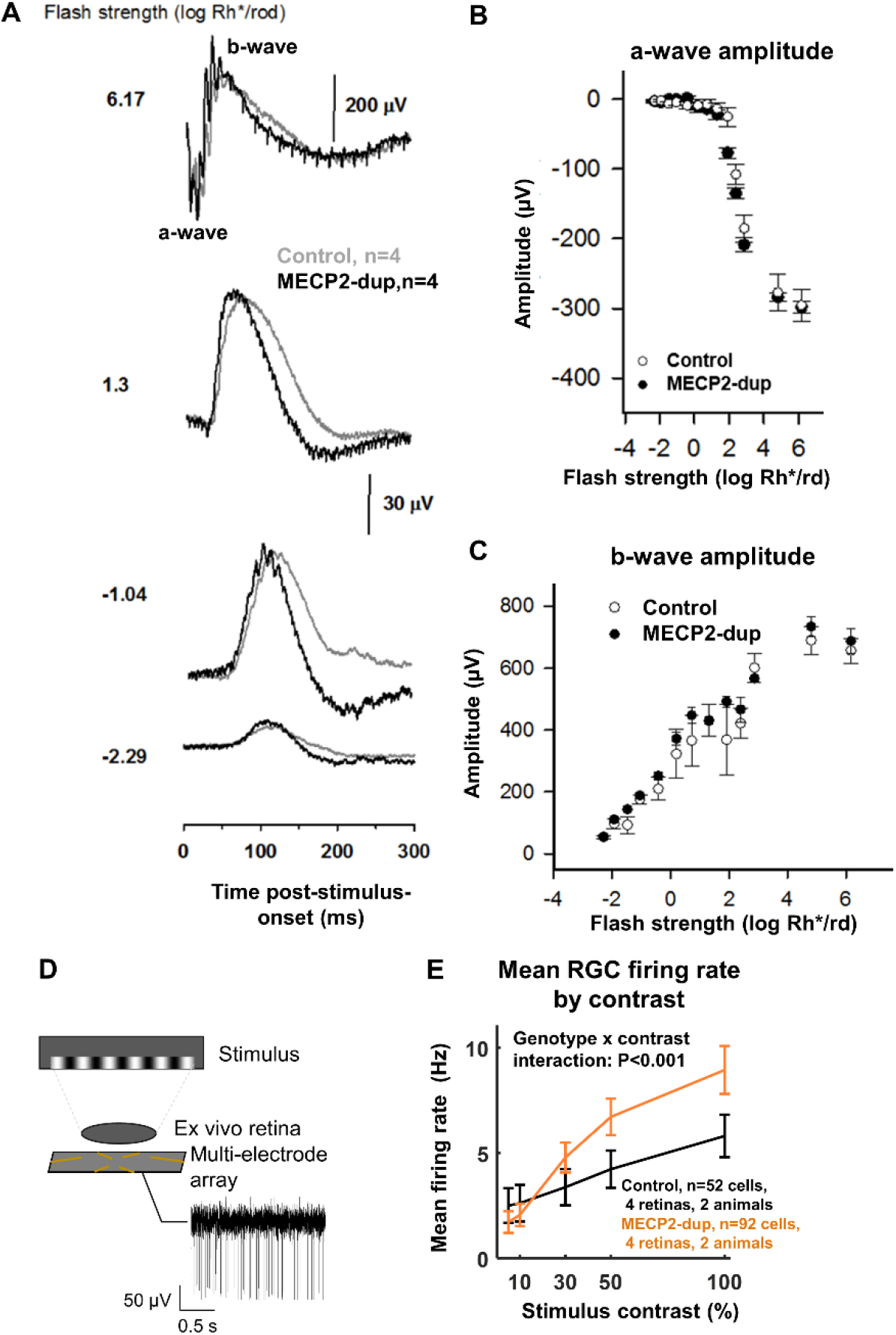
Intact retinal physiology in MECP2-duplication mice. **A.** Mean electroretinogram responses to full field luminance flashes at different flash strengths (numbers at left), recorded from the optic nerve of MECP2-duplication mice and control littermates. Grey, control, n=4 mice. Black: MECP2-duplication, n=4 mice. Axes labeled in figure. **B**. Amplitude of the a-wave (initial negative response, labeled in panel A, produced by depolarization of the photoreceptor layer) as a function of flash strength. **C**. Amplitude of the b-wave (large positive response, labeled in panel A, produced by depolarization in the inner retinal layer) as a function of flash strength. **D**. Schematic illustration of ex vivo retinal ganglion cell (RGC) multielectrode array extracellular recording. The stimulus paradigm consisted in randomly interleaved moving square gratings, 8 directions (45° apart), 5 contrasts (5, 10, 30, 50, and 100%), 20 repetitions, 1 sec stimulus duration, 1.5 sec interstimulus interval (mean luminance background). **E**. Mean RGC firing rate by stimulus contrast. Control littermates: n= 52 cells from 4 retinas, 2 animals. MECP2-duplication: n=95 cells from 4 retinas, 2 animals. P<0.0001, t=-0.12.2, genotype x contrast interaction; P=0.02, t=2.2, effect of genotype; P=0.0005, t=3.4, effect of contrast; n=55 control cells, 97 MECP2-duplication cells, linear mixed-models ANOVA, neurons nested within retinae. RGC firing rates were similar between genotypes at 30% contrast, the contrast at which we see the most dramatic differences in visual cortical neurons (**Fig. 1F**), suggesting that decreased V1 responses at low contrast are not inherited from the retina. The slight increase in high-contrast responses we see in V1 (most clearly seen in **Fig. 1B**) could be due in part to the increased RGC firing rates we see at 100% contrast.

### Possible perceptual effects of abnormal signal-to-noise across contrasts

To begin to evaluate perceptual effects of visual system abnormalities in MECP2-duplication mice, we measured the optokinetic reflex (**Fig. 5**). Animals (n=8 per genotype) were placed head-fixed in a chamber surrounded on 3 sides by monitors displaying horizontally-drifting vertical gratings at different contrasts and spatial frequencies, while pupil location was tracked with an infrared camera (**Fig. 5A**). Similar or greater numbers of eye-tracking movements (ETMs) were observed across spatial frequencies in mutants (**Fig. 5B**), indicating that visual acuity is intact in MECP2-duplication mice. Interestingly, while similar numbers of eye-tracking movements (ETMs) were observed at high contrasts between genotypes (**Fig. 5C**), ETM rate decreased more in control mice at low contrast compared to mutants. This suggests that perceptual sensitivity was increased at low contrasts in mutant animals compared to controls. Our retinal recordings suggest that this does not happen in the retina, but it might be a property of increased response reliability in the subsequent circuits including the visual cortex, tectum, and/or midbrain (Giolli et al., 2006; Liu et al., 2016).

**Fig. 5.**
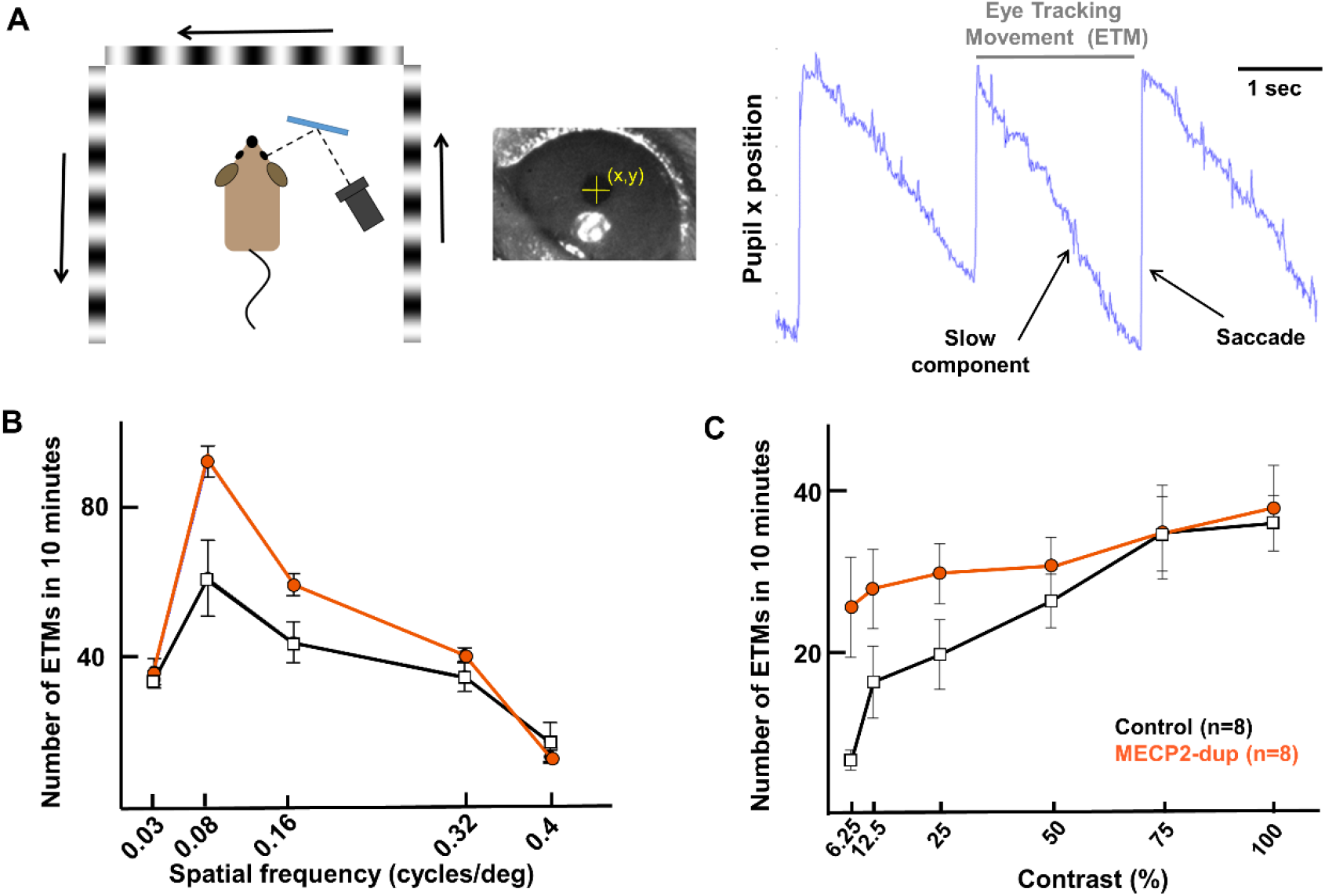
Optokinetic reflex across contrasts and spatial frequencies in MECP2-duplication mice. **A.** *Left panel:* Schematic of optokinetic reflex (OKR) experiment setup. Animals were head-fixed in a chamber surrounded on 3 sides by monitors showing horizontal gratings at different contrasts and spatial frequencies. *Right panel:* The mouse’s pupil was automatically tracked by a custom MATLAB algorithm. OKR movements were parsed into slow eye-tracking movements (ETMs) and fast saccades and counted across 10 minute periods. **B**. Visual acuity assessed by optokinetic nystagmus. Number of ETMs in 10 minutes averaged across animals (n=8 per genotype) at each contrast. Contrast was constant at 100%. Error bars show standard error of the mean. **C**. ETMs reported as in B but as a function of contrast. Spatial frequency was constant at 0.03 cycles per degree. Note the greater fall-off of ETMs in controls compared to mutants at lower contrasts, indicative of enhanced visual sensitivity in MECP2-duplication mice at low contrasts.

## DISCUSSION

We assessed in vivo responses to oriented gratings in layer 2/3 of the primary visual cortex (area V1) of the MECP2-duplication mouse model of autism. MECP2-duplication neurons respond less strongly at low contrasts but similarly at high contrasts compared to littermate controls, leading to more linear contrast-response functions. Interestingly, this does not necessarily imply defective encoding of visual stimuli compared to controls: Mean-normalized response variability, as measured by the Fano factor, drops significantly at low contrasts in mutants compared to controls, and OKR responsiveness is enanced, reflecting an increase in response reliability that could compensate for decreased response amplitude for adequate sensory encoding.

In the primary sensory cortex of normal animals, neuronal responses are typically variable, fluctuating significantly even to identical repetitions of the same stimulus (Werner and Mountcastle, 1963). Neuronal response variability has multiple sources (Faisal et al., 2008; Gjorgjieva et al., 2016; Nolte et al., 2019), but one of the chief contributions arises through the interaction of feedforward sensory-evoked activity with endogenously generated activity fluctuations (Stringer et al., 2019). Endogenous activity manifests as large-scale neural activity modulations across brain areas, mediated in part by feedback projections and reflecting, among others, behaviorally-relevant brain state fluctuations (Ecker et al., 2014; Lin et al., 2015; Okun et al., 2015; Denfield et al., 2017; Stringer et al., 2019), mnemonic activity patterns enabling storage and retrieval of sensory representations (Miller et al., 2014), and spontaneous activity patterns corresponding to Bayesian priors used for statistical inference (Berkes et al., 2011; Orbán et al., 2016). The coupling of neurons to these global patterns of activity varies across individual cells (Okun et al., 2015; Lee et al., 2019), timescales (Okun et al., 2019), and behavioral state (Canolty et al., 2012), making an important contribution to the variability of individual neuronal responses. The increased variability in sensory-evoked responses in autism patients have led to the suggestion that the interaction between sensory evoked and endogenous activity is disrupted in ASD cortex leading to a loss of response reliability (Heeger et al., 2017).

The fact that visual response reliability is higher in MECP2-duplication animals (**Fig. 2,3**) shows that sensory encoding is not universally characterized by increased variability in autism spectrum disorders, as has been previously hypothesized (Dinstein et al., 2012). Indeed, area V1 layer 2/3 neuronal responses fluctuated less from trial to trial in MECP2-duplication animals compared to littermate controls. Two separate mechanisms account for this difference: First, V1 neuropil responses, which provide an LFP-like read-out of large-scale neuronal input activity fluctuations (Kerr et al., 2005; Greenberg et al., 2008; Lee et al., 2019), exhibited decreased variability in mutants (**Fig. 3A**). Such fluctuations are known to drive trial-to-trial variability in V1 neurons (Ecker et al., 2014). Second, mutant neurons exhibited weaker coupling to these endogenous fluctuations, particularly at low contrast (**Fig. 3B,D**). Notably, these mechanisms are unlikely to be inherited from the retina: retinal ganglion cell response amplitudes were not diminished and Fano factors remained unchanged at low contrast (**Fig. 4**). Instead, our data suggest that thalamocortical mechanisms regulating V1 response reliability (Heeger, 1992; Nauhaus et al., 2009) are disrupted in the MECP2-duplication mouse, particularly at low (<~30%) contrasts.

It is interesting to speculate about candidate mechanisms that may explain our observations. Increased tonic inhibitory neuron activity could explain both suppressed response amplitude and decreased variability at low contrasts (Shen et al., 2011; Li et al., 2012; Thiele et al., 2012; Rikhye and Sur, 2015; Zhu et al., 2015). Excessive inhibition has already been demonstrated in MECP2-overexpressing mice (Na et al., 2012, 2014), and GABA antagonism has been shown to rescue phenotypes (Na et al., 2014). For example, increased tone in mutant somatostatin neurons could inhibit dendritic tufts in layer 1 weakening feedback from higher areas, thereby decreasing associated trial-to-trial variability (Rikhye et al., 2017). In contrast, our results do not align well with theories of impaired GABAergic inhibition, which lead to increased excitatory-inhibitory balance in autism (Vattikuti and Chow, 2010). Contrast response functions in mutant mice lay below those of littermate controls, suggesting inhibition mediated normalization mechanisms are, if anything, stronger in mutant animals. Other potential contributing mechanisms include failure of the local recurrent circuitry between L4 and L2/3 pyramidal neurons (Anderson et al., 2000; Nauhaus et al., 2009; Lien and Scanziani, 2013; Cossell et al., 2015) or perhaps the thalamo-cortical loop (Noutel et al., 2011; Yagasaki et al., 2018). Altered intrinsic cellular properties, such as firing thresholds(Anderson et al., 2000), may also contribute, though, to our knowledge such changes have not been described in MECP2-duplication animals (Lu et al., 2016). Future work will be required to further differentiate between these mechanisms.

Optokinetic responses at low contrast were enhanced in mutant animals (**Fig. 5**) and it is tempting to speculate that this may have something to do with the increased reliability we observed at similar contrasts. If so, it may well be that the observed increase in reliability has also other behavioral correlates. Be that as it may, the fact that OKR were preserved in mutants is certainly reassuring, since it excludes the possibility that low visual acuity or other peripheral optical issues could have confounded our observations. Our finding of enhanced contrast-sensitivity by optokinetic reflex measurements is similar to what was seen in the BTBR autism mouse model (Cheng et al., 2020), which also shows dendritic overgrowth and excessive ERK signaling similar to what has been observed in the *MECP2*-duplication mouse (Jiang et al., 2013; Ash et al., 2017).

Interestingly, *decreased* V1 neuron response reliability has been observed in response to natural movie stimulation in the mouse model of Rett syndrome (MeCP2 loss-of-function) (Banerjee et al., 2016). It is perhaps not surprising that this is opposite to our observations in MECP2-duplication mice, as MeCP2 loss vs gain effects have also been observed to be opposite for other neuronal phenotypes, including long term potentiation (Collins et al., 2004; Moretti et al., 2006), synapse strength/number (Chao et al., 2007; Lu et al., 2016), and dendritic spine motility (Landi et al., 2011; Jiang et al., 2013). More generally, there are several conflicting reports arguing for increased (Coskun et al., 2009; Greenaway et al., 2013; Butler et al., 2017) versus decreased (Milne, 2011; Dinstein et al., 2012; Weinger et al., 2014; Haigh et al., 2015; Banerjee et al., 2016; Kovarski et al., 2019) visual response reliability in autism spectrum disorders. The reasons for this have yet to be fully clarified. One proposed explanation hinges on different recording modalities (Butler et al., 2017). For example, the majority of fMRI studies in autism (Haigh et al., 2015) report increased variability. The decreased variability we report here was evident in neuropil fluctuations, which, as a readout of overall population activity, is expected to correlate well with the BOLD (Blood Oxygen Level Dependent) signal (Logothetis et al., 2001; Kerr et al., 2005). However, the possibility remains that BOLD, which has a vascular origin, may partially reflect variability in vascular tone (Hillman, 2014) that may also be differentially affected in autism (Reynell and Harris, 2013). We note in this regard that visual-evoked field potentials, which correlate more closely to neuropil activity (Kerr et al., 2005), have demonstrated normal or increased reliability in autism (Coskun et al., 2009; Butler et al., 2017). Another explanation relates to experimental differences in attentional factors and brain state. In (Dinstein et al., 2012), subjects performed a letter detection task, in (Milne, 2011) and (Weinger et al., 2014) participants were asked to fixate, while in (Kovarski et al., 2019) subjects did not have a task to perform; therefore attentional allocation to the visual stimulus and overall alertness may have fluctuated more over time in ASD subjects vs. controls (Ecker et al., 2014, 2016; Denfield et al., 2017). In our study mice were fentanyl-sedated, so activity fluctuations due to attentional state should be minimal. Finally, perhaps the most likely explanation is that different cortical circuit mechanisms of malfunction operate in different subtypes of autism, even though they result in overall similar core phenotypes (Ramocki and Zoghbi, 2008; Lu et al., 2016; Ash et al., 2021).

One limitation of our study is that recordings were made under fentanyl-dexmetotomidine sedation. Under this form of sedation, animals remain awake but in a state of heavy physiological relaxation (Souter et al., 2007). This regimen tends to generate responses that are closer to quiet wakefulness compared to full anesthesia with ketamine or gas anesthetics (Mrsic-Flogel et al., 2007; Lee et al., 2017). Assessing response reliability under awake behaving conditions is a key area of future work, which will require precise monitoring of the behavioral state (Stringer et al., 2019).

In summary, we report that endogenous activity is less variable and that neuronal coupling to global patterns of activity is weaker in MECP2-duplication animals compared to littermate controls. These results point out a potential mechanism for the “hypo-prior” theory of autism (Pellicano and Burr, 2012). In this theory, Bayesian priors are communicated to local circuits through feedback connections and projections from other areas (Ma et al., 2006; Berkes et al., 2011). Since V1 neurons in mutant animals are weakly modulated by fluctuations of endogenous activity during sensory processing, they may be less able to properly process Bayesian priors embedded in global patterns of activity arising from higher areas. This relative reduction in global information processing (weak coherence theory (Happé, 2021)) could result in more rigid or dominant sensory feature representations in the local circuitry, with potentially detrimental information processing consequences. For example, decreased coupling to endogenously generated activity can produce more reliable but less flexible neural representations, potentially leading to enhanced perceptual sensitivity for simple features but impaired capacity for global computations that require coordination across areas (Robertson and Baron-Cohen, 2017). The failure of neuronal stochastic behavior may also result in poor cortical circuit regularization during learning, leading to excessively rigid neuronal network architectures and failure of generalization (Ash et al., 2017, 2021). These effects may be compounded over time, potentially contributing to stereotyped patterns of circuit activity, repetitive patterns of behavior and behavioral inflexibility (Harris et al., 2015). It remains an appealing hypothesis that abnormal balance between circuit reliability and stochasticity could lead to neural circuit inflexibility that may underlie behavioral inflexibility in autism spectrum disorders (Ash et al., 2021).

## METHODS

### Experimental procedures

#### Animals

All experiments and animal procedures were performed in accordance with guidelines of the National Institutes of Health for the care and use of laboratory animals and were approved by the IACUC at Baylor College of Medicine and Brigham and Women’s Hospital (BWH) Institution Animal Care and Use Committee. In the main experiment (Fig. 1-4F), C57Bl6J mice were crossed to FVB MECP2-duplication (*Tg1*, Collins et al., 2004) mice to generate F1 C57;FVB MECP2-duplication mice and nontransgenic littermate controls. In the replication experiment (Fig. 3), C57Bl6j Thy1-GCaMP6S mice were crossed to 129-background MECP2-duplication mice to generate F1 129;FVB MECP2-duplication;Thy1-GCaMP6 mice and thy1-GCaMP6 littermate controls. Experiments were performed in 4-6-month-old animals.

#### Surgery

All procedures were carried out according to animal welfare guidelines authorized by the Baylor College of Medicine and Brigham and Women’s Hospital IACUC committees. Surgeries were performed following (Holtmaat et al., 2009) and (Mostany and Portera-Cailliau, 2011). Mice were anesthetized with 1.5% isoflurane. The mouse head was fixed in a stereotactic stage (Kopf Instruments), and eyes were protected with a thin layer of polydimethylsiloxane (30,000 cst, Sigma-Aldrich). The scalp was shaved and disinfected with chlorhexidine, and the scalp was resected. A 3 mm wide circular craniotomy centered 2.5 mm lateral of the lambda suture was made, targeting the middle of the monocular region of left V1 (Chen et al., 2013). Fifty nL aliquots of AAV-syn-GCaMP6s virus (1:10 dilution) were injected at 2-3 sites ~ 600 microns apart avoiding large blood vessels, targeted to the posterior 2/3 of the window. A glass cover slip was placed into the craniotomy and sealed with cyanoacrylate, krazy glue, and dental cement. A custom-made titanium headplate was attached to the skull with dental acrylic (Lang Dental).

#### Imaging

About 3 weeks following surgery and virus injection, animals’ eyes were visually inspected to exclude mice with cataracts (rare in the C57;FVB F1 background). Mice were sedated with fentanyl (0.5 mg/kg) and dexmetetomidine (0.5 mg/kg) and 20-30 minutes later head-fixed under the microscope. A plane of neurons was selected at 170-200 microns depth, and correct placement of the mouse’s head relative to the visual stimulus was confirmed by a retinotopy stimulus. A 20x, 0.95 NA (Olympus) in a modified Prairie Ultima IV two-photon laser scanning microscope (Prairie Technologies, Middleton, WI), fed by a Chameleon Ultra II laser (Coherent, Santa Clara, CA), was used for 2-photon imaging. Images were acquired by spiral scan at 256×256 micron field of view (FOV), 0.9-1 microns per pixel, 5.5-7 Hz fps. In the replication experiment, images were acquired at 0.92-1.25 microns per pixel, 3.4-4.2 Hz fps. Appropriate imaging sites were screened under 2-photon by recording GCaMP6 responses to a retinotopy stimulus to confirm appropriate virus expression and targeting to V1 (in our hands V1 neurons respond highly reliably to this stimulus).

#### Visual stimulation

Visual stimuli were generated in MATLAB and displayed using Psychtoolbox (Brainard, 1997). The stimuli were presented on an LCD monitor (DELL 2408WFP, Dell, Texas, USA) at 60 Hz frame rate, positioned 32 cm in front of the right eye, centered at 45° clockwise from the mouse’s body axis. The visual angle of the screen spanned 540 elevation and 780 azimuth. The screen was gamma-corrected, and the mean luminance level was photopic at 80 cd/m^2^. Our visual stimulation paradigm consisted of square-wave drifting grating stimuli. Twelve different grating orientations (30°, 60°,… 360°) at 3 contrasts (10%, 30%, 100% Michelson contrast (Michelson, 1927) were shown, giving a total of 36 stimuli pseudorandomly interleaved. Temporal frequency was set to 2Hz, and spatial frequency was set to 0.04 cycles/degree. The duration of each trial was 2 seconds on, 2 seconds off. Trials were presented in 2 blocks of 5-6 repetitions per stimulus orientation/contrast, giving a total of 10-12 repetitions per stimulus orientation/contrast. Each grating stimulus was followed by an inter-stimulus interval during which a full-field gray screen at the same mean luminance (background illumination) was presented.

#### Electroretinograms

ERGs were performed as in (Tse et al., 2015). Briefly, mice were dark-adapted overnight. Under dim red light, mice were anesthetized with ketamine-xylazine (100/10 mg/ml), and their pupils were dilated with a single drop of 0.5% mydryacil and 2.5% phenylephrine. A small amount of 2.5% methylcellulose gel was placed on the eye and a platinum electrode was placed in contact with the center of the cornea. Reference and ground electrodes were placed on the forehead and tail respectively. Mice were placed in a Ganzfeld dome coated with reflective white paint. Scotopic a-wave and b-wave measurements were made with 0.5 ms square flashes of 503 nm peak wavelength, 40 repetitions, 2 seconds between each stimulus. Electrical signals were analyzed with custom software in MATLAB. The b-wave was digitally filtered using the *filtfilt* function in MATLAB (low-pass filter, Fc=60 Hz) to remove oscillatory potentials. The a-wave was measured from baseline to the trough of the initial negative deflection (unfiltered) and the b-wave was measured from the a-wave trough to the peak of the subsequent positive deflection (filtered).

#### Retinal ganglion cell recording

Experimental preparation was performed as in (Cowan et al., 2016; Sabharwal et al., 2017; Seilheimer et al., 2020). Two 16-week-old MECP2-duplication and two litter-mate controls were used in experiments. Prior to euthanization, mice were dark adapted for at least 90 min. Eyes were removed under infrared illumination using night vision (Nitemare, BE Meyers, Oregon) and their retinas were dissected in a dish containing carboxygenated recording solution. Retinas were placed ganglion cell side up onto nitrocellulose filter paper (0.45 μm HA, Millipore) and transferred onto an electrode array where the preparation was retained with a plastic ring.

The retina was kept at 35.6°C and perfused at 2 mL/min with prewarmed and carboxygenated (95% O_2_, 5% CO_2_) recording medium (in mmol/L: NaCl, 124; KCl, 2.5; CaCl_2_, 2; MgCl_2_, 2; NaH_2_PO_4_, 1.25; NaHCO_3_, 26; and glucose, 22) at pH 7.35. The multielectrode array (MEA-60, Multichannel Systems, Tübingen Germany) had 60 electrodes spaced 100 μm apart and with diameters of 10 μm. Ganglion cell action potentials were recorded at 20 KHz and prefiltered with a 0.1 Hz high-pass hardware filter. Visual stimuli were presented from a computer monitor (Dell, SXGA-JF311-5100) optically reduced and presented from below the MEA onto the ganglion cell side of the retina. The stimulus paradigm consisted in randomly interleaved moving square gratings, 8 directions (45° apart), 5 contrasts (5, 10, 30, 50, and 100%), 20 repetitions, 1 sec stimulus duration, 1.5 sec interstimulus interval (mean luminance background).

#### Visual acuity assessment by optokinetic reflex

Nystagmoid responses to square-wave drifting gratings were recorded in 8 MECP2-duplication mice and 8 littermate controls. The stimulus was presented on three screens, positioned around the mouse to cover ~270° of mouse’s visual field. The center of each screen was located at 27 cm from the mouse. An infrared camera (GC660, Allied Visual Technologies) was used to record the movements of the right eye at 60Hz. Ten-to twenty-minute-long movies were analyzed off-line using custom Matlab code to determine the position of the pupil relative to the position of the reflection of the infrared light source on the surface of the cornea. Periods containing eyeblink artifacts were removed. Horizontal eye tracking movements (ETMs), consisting of slow pursuit movement in the direction of the global stimulus drift ±89° and preceded by a saccade in the opposite direction ±89°, were selected for further analysis. A linear fit was applied to the slow pursuit phase of each EM, and EMs with r^2^ > 0.5 were accepted for further analysis. The total number of ETMs in 10 minutes was quantified for each contrast and spatial frequency.

### Data analysis

#### Preprocessing

Movies were motion-corrected using a Fourier transform method (Yatsenko et al., 2015). For cell identification, a local region-of-interest (ROI) over a cell body was manually defined with a circular disk to cover the cell body, then scanned for the pixel with the highest fluorescence value within the disk (cell body center) using the matlab function roigui from Chen et al., 2013. The boundary of the region containing the cell signal was then defined by thresholding at 0.5x maximum fluorescence within the disk along the polar coordinates. To correct for slow signal drift over time, the signal time series from each pixel was high-pass filtered (HPF) at 0.05 Hz using the discrete cosine transformation. We then measured the ratio between the mean calcium signal within the lumen of non-radial blood vessels (≤10 μm) and the surrounding neuropil patch (Kerlin et al., 2010). This gave us an approximate measure of neuropil contamination at the cell soma, the so called contamination scale, whose typical value was 0.6. To correct for the neuropil contamination at the soma, the mean fluorescence of the adjacent neuropil patch, *Fn*, was subtracted using the contamination scale *S: F_correct_ = F - S*Fn*. The patch *Fn* formed an annulus with a radius of 7-15 μm centered around the soma, excluding pixels that belonged to other cell bodies. In the calcium images, the majority of optical contamination was particularly along the z-axis, and local neuropil activity was highly coherent. The correction factor compensated for neuropil contamination along the z-axis. The contamination of the neuronal signal by the neuropil signal varied with cell size as well as with the in-plane diameter of the soma cross-section. We therefore examined a range of values around the empirically estimated value *S* was in range 0.5-0.6 and verified that reasonable variation in the level of contamination (*S*) did not significantly affect our conclusions. Note that, unless explicitly mentioned, all cell responses shown were calculated by implementing a neuropil contamination correction with the empirically determined factor *S* = 0.6. All the analyses presented here are based on the calcium fluorescence dF/F signal per recording.

#### Cell and Neuropil Analysis

To calculate percent fluorescent change in a cell, the mean dF/*F* was computed by projecting the dF/F matrix (time-point samples x pixels in the cell) onto the spatial filter *a* (pixels x1) and normalizing by the sum of the coefficients *a,* to produce a weighted mean. The filter *a_i_* was optimized using deconvolution. Around each cell (ROI defined as in preprocessing section above), a local neuropil patch was defined by selecting an annulus centered on the middle of the cell body. A variety of inner outer radii of the annulus were chosen. The annulus sizes ranged from 7-9 μm, 9-11 μm, and so on by incrementally increasing the radius size with a step size of 2μm, and also one for a large patch size: 7 μm to 15 μm, respectively. We chose the minimum inner radius as 7 μm to minimize contamination from the cell signal onto the neuropil patch. In accordance with a previous report for the size of cell somata in mouse visual cortex (Braitenberg and Schüz, 1998), this was indeed large enough to exclude any pixels from other cell somata, by visual inspection. Other cell bodies, glia, and dark blood vessel regions were similarly excluded. After the application of high-pass filtering (cut-off at 0.05 Hz), the dF/F was calculated in single pixels with the same method as for the cell somata, and then averaged across all pixels within the patch.

#### Spontaneous activity

Five minutes of spontaneous activity were recorded in each animal at the beginning of experiments. dF/F traces were thresholded with all dF/F <0.8 set to 0 as in (Lu et al., 2016), and peaks were detected using automated scripts in MATLAB, to quantify spontaneous event rate and event amplitude. Similar results were scene for a range of dF/F thresholds (0.2 to 1).

#### Responsive cells

Cell responsiveness was based on orientation selectivity recordings. Visually responsive cells were selected if there was a significant difference in mean dF/F plus standard deviation around the peak of the expected response (i.e. 12 frames after stimulus onset ±8 frames; total number of frames during stimulus=22 frames) compared to the pre-stimulus response.

#### Calculation of response amplitude

The mean response to the visual stimulus was calculated for each ROI (cell or neuropil) as illustrated in Fig. 1C. Each trial response was calculated as the average of the 3 frames centered at the peak of the mean 100%-contrast response across all orientations +/- 1 frame, minus the baseline activity (the average of first 3 frames post-stimulus-onset). These frames were used for calculating the baseline activity on the observation that in our data the GCaMP6s calcium signal generally reached its minimum at these frames. Mean response amplitude was then calculated as the average across all trials for each stimulus orientation (independent of direction).

#### Analysis of tuning properties and response reliability

Orientation selectivity index (OSI) was determined using Mazurek et al.’s method (Mazurek et al., 2014) as the normalized measures of “peak to trough” orientation selectivity and direction selectivity: *OSI* = (*R_preferred_ - R_orthogonal_*)/*R_preferred_*. The response at the preferred orientation *R_preferred_* was taken to be the best response to one of the stimulus orientations by measuring the responses at stimulus directions *ϑ*_1_, *ϑ*_2_,…, *ϑ_n_*, then we choose the response at the best *ϑ_i_*,. An OSI value of 0 indicated the absence of orientation selectivity, whereas a value of 1 indicates maximum orientation selectivity. Tuning width was calculated as the full width of the tuning curve at half the mean response to the preferred stimulus. Direction selectivity index (DSI) was calculated as the response to the preferred stimulus minus the response to the null (180-degree-opposite) stimulus, DSI = (*R_preferred_* – *R_null_*)/*R_preferred_*. Fano factor was calculated as the variance in the response amplitude across trials for the preferred or non-preferred stimulus divided by the mean response amplitude for that stimulus, σ^2^/μ.

#### Statistics

Calcium imaging data was analyzed by 2-way ANOVA, followed by Tukey test for multiple comparisons. Retinal recordings were analyzed with the linear mixed-effects models ANOVA, with neurons nested within retinae, nested within animals.

## AUTHOR CONTRIBUTIONS

RTA and SMS conceived of the project and designed experiments, RTA, JP, GP, and SL performed in vivo 2-photon imaging experiments, JAF, RTA, and GP analyzed the in vivo 2-photon imaging data, DL and MS segmented 2-photon images for the replication experiment, ZHT, EF, and AT provided technical and theoretical assistance with experiments, RS and JS performed the ex vivo RGC recordings and analyzed the data, FR performed the optokinetic reflex experiments and analyzed the data, RTA, JAF, and SMS wrote the manuscript.

## ACKNOWLEDGEMENTS

Thanks to H.Y. Zoghbi for advice on experiment design and manuscript preparation, and use of the Tg1 mouse line.

